# DeepEnzyme: a robust deep learning model for improved enzyme turnover number prediction by utilizing features of protein 3D structures

**DOI:** 10.1101/2023.12.09.570923

**Authors:** Tong Wang, Guangming Xiang, Siwei He, Liyun Su, Xuefeng Yan, Hongzhong Lu

## Abstract

Turnover numbers (kcat), which indicate an enzyme’s catalytic efficiency, have a wide range of applications in fields including protein engineering and synthetic biology. Experimentally measuring the enzymes’ kcat is always time-consuming. Recently, the prediction of kcat using deep learning models has mitigated this problem. However, the accuracy and robustness in kcat prediction still needs to be improved significantly, particularly when dealing with enzymes with low sequence similarity compared to those within the training dataset. Herein, we present DeepEnzyme, a cutting-edge deep learning model that combines the most recent Transformer and Graph Convolutional Network (GCN) architectures. To improve the prediction accuracy, DeepEnzyme was trained by leveraging the integrated features from both sequences and 3D structures. Consequently, our model exhibits remarkable robustness when processing enzymes with low sequence similarity compared to those in the training dataset by utilizing additional features from high-quality protein 3D structures. DeepEnzyme also makes it possible to evaluate how point mutations affect the catalytic activity of the enzyme, which helps identify residue sites that are crucial for the catalytic function. In summary, DeepEnzyme represents a pioneering effort in predicting enzymes’ kcat values with superior accuracy and robustness compared to previous algorithms. This advancement will significantly contribute to our comprehension of enzyme function and its evolutionary patterns across species.

## Introduction

The enzyme turnover number (*k*_cat_) represents the maximum number of substrate molecules that an enzyme can convert into a product per unit time under saturating conditions^1,2^. Presently, accurately predicting *k*_cat_ has become imperative for various applications, including protein engineering and enzyme design^3^. Additionally, estimating enzyme *k*_cat_ values at genome-scale is crucial for constructing sophisticated metabolic models to capture the correlations between genotypes and phenotypes^4,5^. Despite the abundant *k*_cat_ values available in the BRENDA^6^ and SABIO-RK^7^ databases, the total number of enzymes with experimentally determined *k*_cat_ remains substantially smaller than the vast number of all sequenced proteins. For instance, less than 1% of the enzymes listed in the UniProt database possess experimentally ascertained *k*_cat_ values^8^.

Recently, it witnessed the breakthroughs in development of various advanced deep learning models, which make it possible to predict protein structure^9^, protein function^10^, and gene expression level^11^ only from primary sequence features. As for genome-scale *k*_cat_ prediction, Heckmann et al.^12^ firstly utilized machine learning models, i.e. the linear elastic net, the decision-tree-based random forest model, to infer the enzyme *k*_cat_ for the model organism *Escherichia coli*. They discovered a correlation between *k*_cat_ and specific enzyme structural features. While those classical machine learning models could successfully predict *k*_cat_ parameters at the genome-scale^12^, its application is only suitable for *E. coli*. Subsequently, the deep learning model-DLKcat, proposed by Li et al.^13^, significantly broadens the scope of *k*_cat_ prediction for nearly all sequenced enzymes with catalyzed substrates. By incorporating features from both the protein sequence and its substrate, DLKcat facilitated high-throughput prediction of enzyme *k*_cat_ and the automatic generation of genome-scale *k*_cat_ profiles. To further enhance the performances of DLkcat, TurNuP integrated comprehensive biochemical reaction information, encompassing fingerprint features from substrates and products, and effectively predicted the *k*_cat_ of wild-type enzymes^14^. It outperforms DLKcat in predicting *k*_cat_ for enzymes with low sequence similarity compared to those in the training dataset. However, there remains room for improvement in the prediction accuracy of TurNuP, given that its R^2^ in test datasets is approximately 0.42, indicating that only a small portion of the variance in *k*_cat_ prediction can be explained by this model.

To a larger extent, the function of a protein is determined by its 3D structure^15^. The structural data provide crucial insights into the spatial arrangement of pivotal functional residues and the interactions between the enzyme and its substrate^16,17^. The composition of amino acids and the distinct folding pattern within the protein structure can significantly influence the accessibility of active sites, substrate binding sites, and the overall stability of the enzyme^18^. These factors collectively impact enzyme function and its catalytic efficiency. Consequently, features derived from protein 3D structures have garnered special attention in the development of advanced deep learning models. These sophisticated models can now accurately predict how amino acid mutations affect the thermal stability of enzymes^19^, as well as identifying the binding sites of ligands on the 3D structure of proteins^20^. Recent breakthroughs in protein structure prediction, exemplified by the release of AlphaFold2^9^, RoseTTAFold^21^, and ColabFold^22^, have substantially reduced the cost in acquiring high-quality protein structures. Thus, it is feasible to build large-scale structure datasets for the training of advanced deep learning models. However, despite this progress, the invaluable protein 3D structural information spanning various species has not been systematically utilized in enzyme *k*_cat_ prediction.

Here, we present a novel deep learning model, named as DeepEnzyme, to enhance the prediction accuracy of enzyme *k*_cat_ based on protein 3D structures. DeepEnzyme combines Graph Convolutional Networks (GCN)^23^, a proven approach for extracting structural properties from proteins^24,25^, with Transformers^26^, which have demonstrated exceptional performance in protein language modeling^27,28^. This fusion allows DeepEnzyme to extract both structure and sequence features from enzymes and substrates. Compared with existing models, DeepEnzyme exhibits superior accuracy and robustness in predicting *k*_cat_ values. Therefore, DeepEnzyme has the potential to expedite the functional analysis of previously unstudied enzymes and provide opportunities for the rational optimization of enzyme activities in conjunction with advanced molecular technologies.

## Results

### Framework of a novel deep learning model for predicting *k*_cat_

The novel deep learning model for *k*_cat_ prediction, DeepEnzyme, was developed by integrating three layers of features extracted respectively from protein 1D sequence, 3D structure, and the related substrate, utilizing distinct computation modules (Fig. 1). Specifically, an excellent Transformer model was employed to extract features from the protein 1D sequence. Simultaneously, each protein 3D structure was transformed as a residue contact map falling within a standard criteria (*C*_*a*_ − *C*_*a*_ < 10Å , Methods). Subsequently, GCN was applied to extract structural features from the protein 3D structures based on the contact map. Concurrently, the RDKit software^29^ was used to extract essential information from the SMILES formulas of the examined substrates, such as fingerprints and adjacency matrices, which were then used as input for a dedicated GCN module to extract the detailed features of substrates. Following that, all feature vectors derived from both enzyme and substrate were merged. This procedure resulted in the construction of a comprehensive embedding vector that encapsulated enzyme and substrate features. Finally, a neural attention approach based on the representation vectors was used to predict the *k*_cat_ value. Using this attention method, the model was able to highlight important hidden features during the prediction of *k*_cat_ values.

**Fig. 1.**
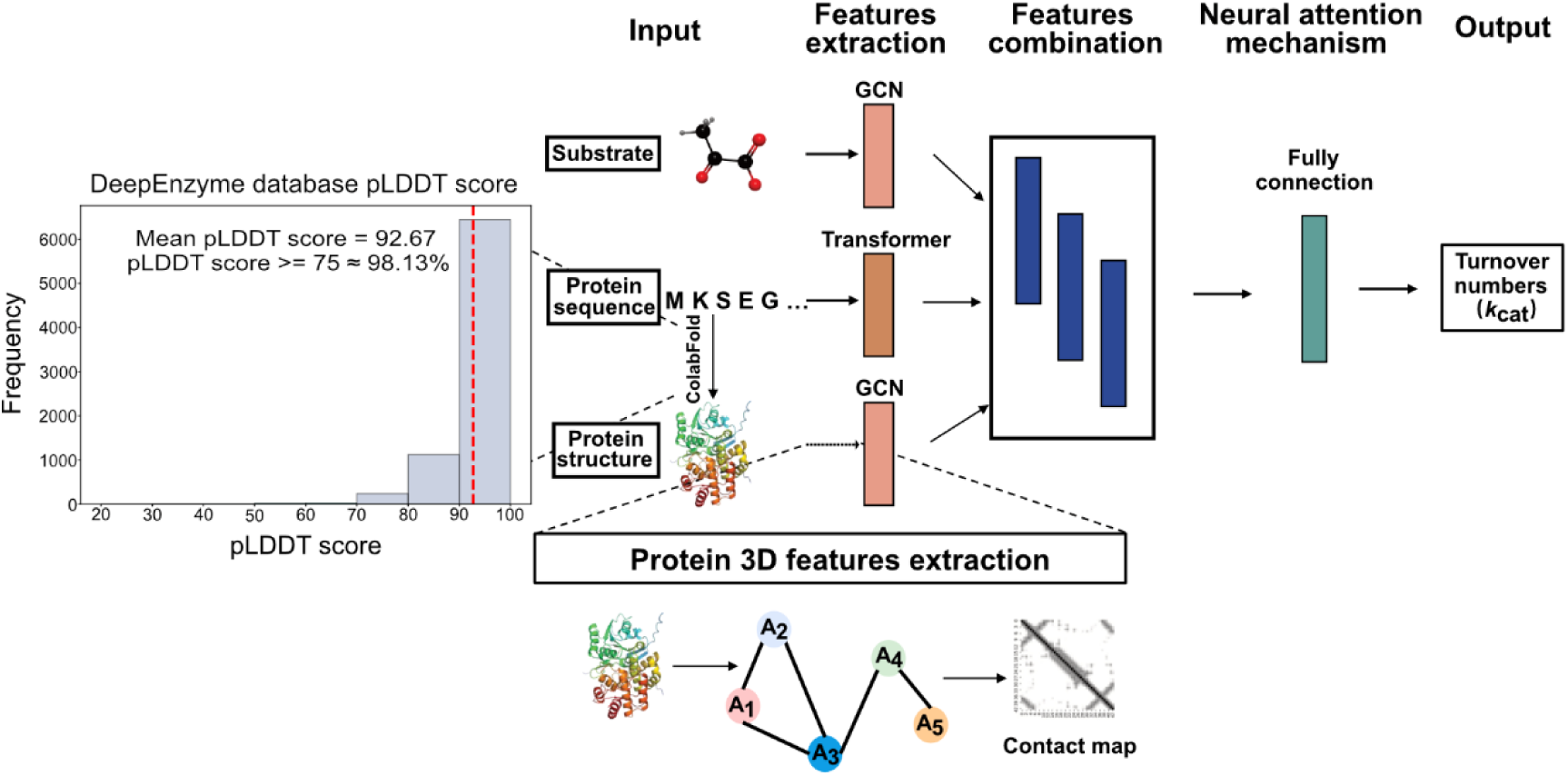
The framework of DeepEnzyme for *k*_cat_ prediction. DeepEnzyme integrates Transformer and GCN models to distill features from both the enzyme and substrate for predicting *k*_cat_. Here, GCN is employed to extract structural features based on protein 3D structures and substrate adjacency matrices; Transformer is utilized to extract sequence features from protein sequences. ColabFold, a protein structure prediction tool, is employed to predict protein 3D structure. The mean pLDDT score for all protein 3D structures used in this work is 92.67, with 98% of the predicted structures achieving a pLDDT score greater than or equal to 75.

Since a significant portion of enzymes in our dataset lacked experimentally determined 3D structures, ColabFold^22^ was utilized to predict the structures for all these proteins. The average predicted local distance difference test (pLDDT) score for all predicted protein 3D structures in this study was 92.67 (Fig. 1), guaranteeing that the structures used for our model training are of high-quality.

### DeepEnzyme exhibits improved performance in *k*_cat_ prediction when taking protein 3D structures into account

Previously, it was demonstrated that the great similarity in protein sequences between the training, validation, and testing datasets can lead to overfitting and poor generalization of deep learning models in *k*_cat_ value prediction^14^. To mitigate this challenge, we preprocessed the data (Methods) to exclude highly similar sequences existing in the original DLKcat dataset^14,30^. Consequently, out of the 16,838 distinct enzyme-substrate pairs, we carefully selected and retained 11,923 of all pairs, enabling us to reconstruct a dataset that could, to some extent, avoid the too high similarity in protein sequences used for model training. This dataset was then randomly partitioned into training, validation, and testing datasets with a ratio at 80%, 10%, and 10%, respectively. This preprocess of the enzymatic dataset allowed us to train and evaluate DeepEnzyme effectively. By mitigating the issue of sequence similarity and incorporating additional protein structure information, DeepEnzyme achieved a high R^2^ value, close to 0.60 (Fig. 2a) on the test dataset after the dedicated training phase, indicating that the model can explain near 60% of the variance in the prediction. Furthermore, the model’s Root Mean Square Error (RMSE) of 0.95 indicates that it has a comparatively low prediction error.

**Fig. 2.**
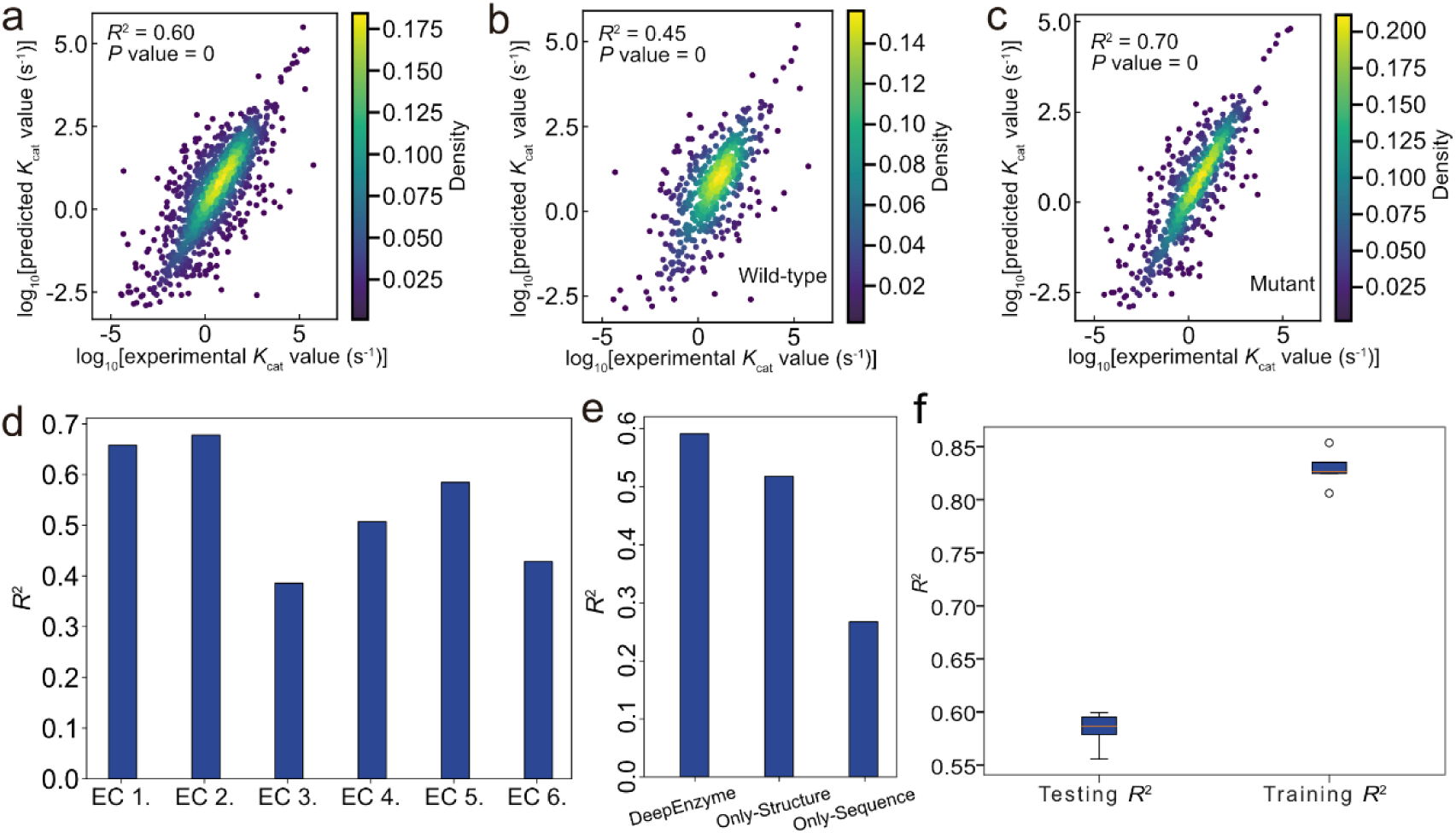
Evaluation of DeepEnzyme performance in *k*_cat_ prediction. (a) The performance of DeepEnzyme on the test dataset was evaluated by the R^2^ and *P* values calculated from the predicted and experimental *k*_cat_ values. (b-c) DeepEnzyme prediction performance on wild-type (b) and mutant (c) enzymes in the test dataset. (d) DeepEnzyme prediction performance for enzymes classified by different EC numbers in the test dataset. (e) The performance of the model with different types of datasets as input. DeepEnzyme: enzyme sequence, enzyme structure, and substrate information are used as inputs for deep learning model training and testing; Only-structure: substrate information and enzyme structure information as inputs; Only-sequence: substrate and enzyme sequence as inputs. In (a)-(e) specific seed was adopted during the calculation. (f) Comparison in average coefficient of determination (R^2^) values for testing dataset and training dataset from 5 rounds of training.

To systematically evaluate the performance of DeepEnzyme, the predicted and measured *k_cat_* values for enzymes from diverse sources and classifications were compared thoroughly. Initially, enzymes were categorized into two main types: wild-type and mutant, based on their records in the public databases. The results indicated that, for mutant enzymes, the R^2^ value was 0.70 (Fig. 2c), whereas for wild-type enzymes, it was only 0.45 (Fig. 2b). Notably, mutant enzymes constituted approximately 59% of all enzymes, suggesting that the prediction accuracy might be influenced by the quantity of enzymes from each group. Next, all enzymes were classified based on the first digit originated from the corresponding EC number. We found that DeepEnzyme exhibits distinct predictive capabilities for enzymes of varying EC numbers. For example, the R^2^ for enzymes from the EC 1 group and the EC 2 group was notably higher (Fig. 2d). This observation is consistent with the fact that the percentages of enzymes from these two groups in the dataset (37%, 26%) are significantly higher than the other enzyme groups, underscoring that the number of enzymes in each group could affect the model prediction capability. Furthermore, to illustrate the enhancement in *k_cat_* prediction achieved by incorporating protein structure, we compared the performance of DeepEnzyme using different types of datasets as input (Fig. 2e). The findings clearly showed that prediction accuracy was much increased by the addition of structural data. Lastly, to avoid the bias in the dataset sampling, the average R^2^ value on the testing dataset from 5 rounds of training was calculated, which achieved at 0.58 by DeepEnzyme, indicating the robustness of model prediction when using different test datasets (Fig. 2f). In summary, DeepEnzyme performs well in *k_cat_* prediction as a whole when taking features of protein 3D structures into account though in some cases the prediction accuracy could be negatively affected by the related data size.

### DeepEnzyme has better performance in *k*_cat_ prediction compared with existing models

To comprehensively compare DeepEnzyme with existing *k*_cat_ prediction tools, two of the latest publicly available models with accessible scripts, TurNuP^14^ and DLKcat^13^, were selected and evaluated together with DeepEnzyme. At the first glance, when only utilizing protein structure and substrate as inputs, DeepEnzyme achieved an R^2^ value of about 0.52 on the test dataset, representing a 30% and 21% improvement compared to TurNuP and DLKcat, respectively (Fig. 3a). Moreover, when utilizing protein sequence, 3D structure and substrate as input, DeepEnzyme exhibited significantly higher prediction accuracy (R^2^ ≈ 0.6) compared to TurNuP and DLKcat. In addition, the RMSE value of DeepEnzyme was 0.95, which was lower than that of DLKcat but slightly higher than TurNuP (Fig. 3b). This discrepancy in RMSE value could be due to TurNuP’s training method, in which the authors omitted the enzyme-catalyzed reactions with exceptionally low or high *k*_cat_ values, resulting in a drop in the model’s final RMSE value^14^. Overall, these results underscore that DeepEnzyme performs well in *k*_cat_ prediction when utilizing protein structure as input, stressing the importance of structure-derived characteristics in accurate *k*_cat_ prediction.

**Fig. 3.**
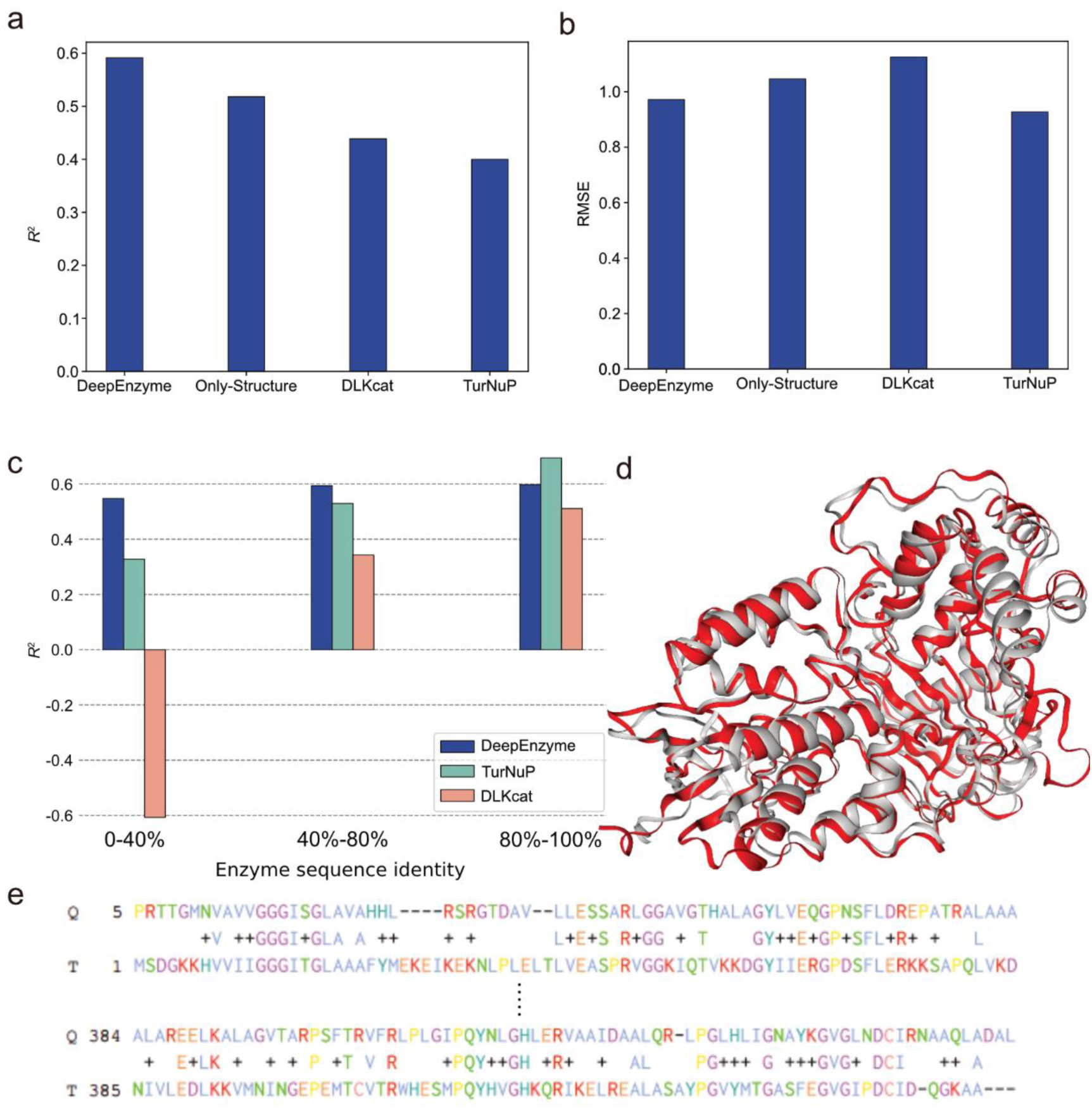
Improved performances of DeepEnzyme in *k*_cat_ prediction compared to existing models, even for protein sequences in the test dataset exhibiting lower similarity compared to those in the training dataset. (a) Comparison of R^2^ values on the test dataset for different models. (b) Comparison of RMSE values on the test dataset for different models. (c) Comparison of R^2^ in *k*_cat_ value prediction for enzymes in the test dataset at different levels of sequence similarity by DeepEnzyme, TurNuP, and DLKcat. (d) Two enzymes from *Myxococcus xanthus* and *Bacillus subtilis*, both with EC numbers 1.3.3.4, are highly similar in protein 3D structure (TM-Score = 0.8762), gray for enzyme from *Myxococcus xanthus* and red for enzyme from *Bacillus subtilis*. (e) The similarity of the amino acid sequences for the above two enzymes is 27% (Q for enzyme from *Myxococcus xanthus*, T for enzyme from *Bacillus subtilis*).

Sequence similarity has a significant impact on deep learning model performance in predicting enzyme function, particularly in predicting the substrates catalyzed by individual enzymes. This impact has been thoroughly explored, as previously discussed by Alexander Kroll et al^30^. Notably, the prediction accuracy of a deep learning model can significantly diminish when the protein sequences in the test dataset exhibit lower similarity to those in the training dataset. To assess the performance of DeepEnzyme at various levels of sequence similarity, we employed the MMseqs2^31^ method to compute sequence similarity between the test and training datasets. Subsequently, based on the calculated similarity, the test dataset was divided into three distinct groups: 0–40%, 40– 80%, and 80–100%. We then calculated the R^2^ value for each group (Fig. 3c). Remarkably, DeepEnzyme showed an impressive R^2^ value of 0.55 even when the sequence similarity spanned from 0 to 40%, demonstrating its robustness in *k*_cat_ prediction under low sequence similarity situations.

In a detailed comparison of the predicted performance of TurNuP, DLKcat, and DeepEnzyme on enzymes with varying levels of sequence similarity (ranging from 80–100% to 0–40%), it was observed that TurNuP experienced a substantial decrease in R^2^, whereas the corresponding R^2^ remained relatively stable for DeepEnzyme (Fig. 3c). It may be due to the fact that, during model training phase, DeepEnzyme learnt useful features of protein 3D structures , which are closely correlated to the enzyme function. In reality, homologous enzymes can exhibit higher conservation in 3D structures while displaying divergent evolutionary paths in terms of sequences. As one of the typical examples, there are two protein sequences in our datasets having the same EC number -1.3.3.4, the final common enzyme in the chlorophyll and heme biosynthesis, from different species (*Myxococcus xanthus* and *Bacillus subtilis*). Calculated by Foldseek and US-align^32,33^, it revealed that the structural similarity between those two enzymes is as high as 0.88 (Fig. 3d), although their sequence similarity is only 0.27 (Fig. 3e). Together, our findings demonstrate DeepEnzyme’s capacity to successfully use 3D structural information for enhanced prediction, while also showcasing robustness in prediction accuracy over a wide range of sequence similarities.

### Evaluation of DeepEnzyme’s prediction capability using enzyme saturation mutagenesis datasets

To validate the capability of DeepEnzyme in predicting the impact of mutations on the catalytic efficiency of enzymes, we firstly compared the predicted *k*_cat_ from our model with the high-throughput experimental datasets for CYP2C9, a critical enzyme in drug metabolism and personalized therapy^34–37^. Based on the enzyme activity^37^, all variants of CYP2C9 were categorized into three groups: missense variants, synonymous variants, and nonsense variants. According to the experimental measurement from Amorosi et al.^37^, the nonsense variants of CYP2C9 have a relatively lower activity score and cellular protein abundance score compared to other mutations. We utilized the well-trained DeepEnzyme to predict and compare the *k*_cat_ values of missense variants and nonsense variants (no comparison was conducted for synonymous variants as they do not undergo sequence alterations). The result clearly indicates that the predicted *k*_cat_ values for nonsense variants (median: *k*_cat_ =1.60s^-1^) were lower than those of missense variants (median: *k*_cat_ = 1.97s^-1^) (Fig. 4a). Additionally, a significant difference was observed between the two groups (*P* value = 1.6×10^-94^). Interestingly, such a result is consistent with experimental evidence represented by the activity score (Fig. 4b). These results confirm a clear difference in expected *k*_cat_ values between missense and nonsense variants, highlighting the unique capability of DeepEnzyme to detect and discriminate these kinds of variations.

**Fig. 4.**
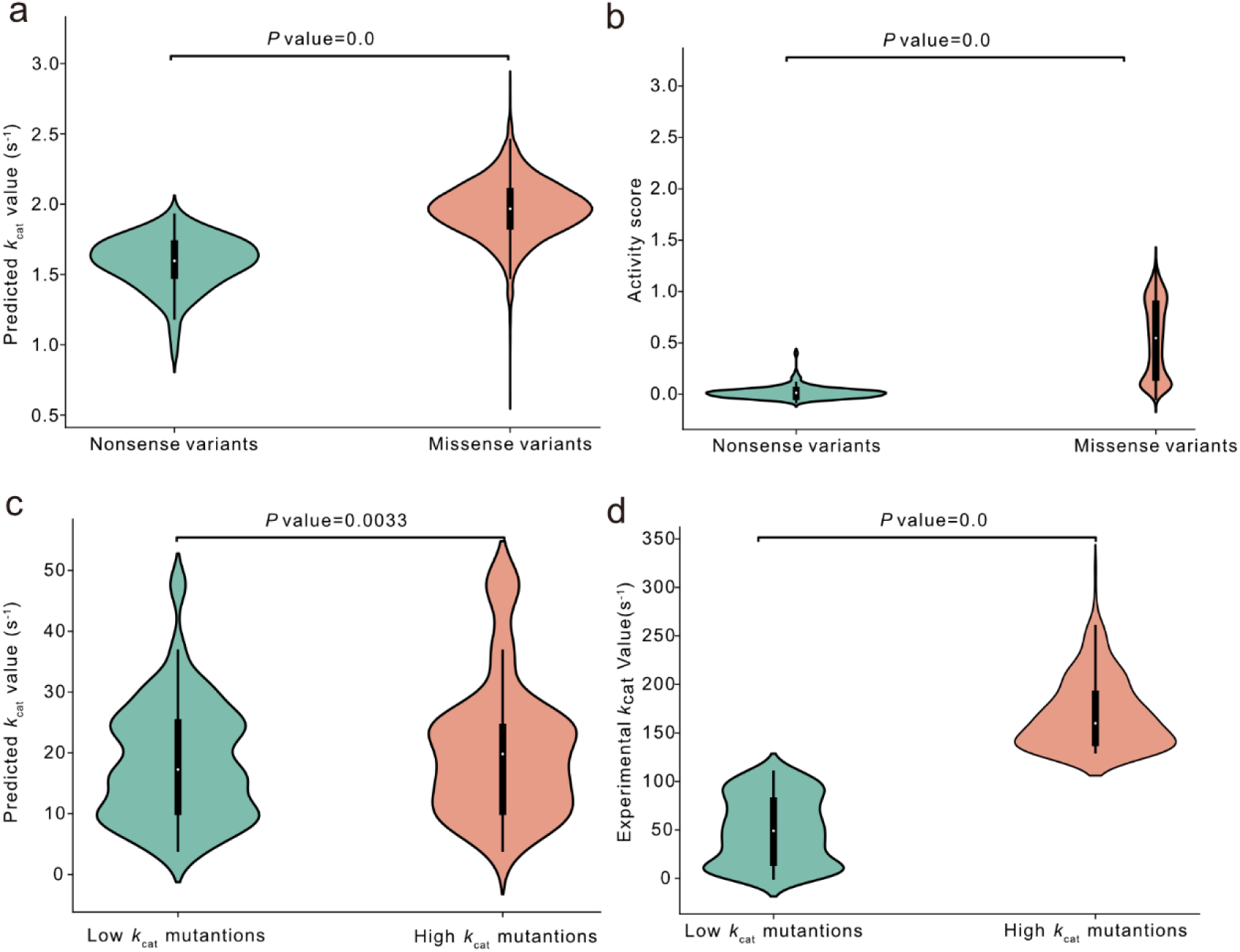
Analysis of the prediction ability of DeepEnzyme for two enzymes with saturation mutagenesis datasets. (a) Comparison of predicted results for different CYP2C9 variants: red for missense variants, green for nonsense variants. (b) Comparison of experimental activity score for different CYP2C9 variants: red for missense variants, green for nonsense variants. In (a) and (b) all variants were divided into three groups: missense variants, synonymous variants, and nonsense variants according to the previous report^37^. (c) Comparison of predicted *k*_cat_ values for different PafA mutations: green for low *k*_cat_ mutations, red for high *k*_cat_ mutations. (d) Comparison of experimentally measured *k*_cat_ values for different PafA mutations: green for low *k*_cat_ mutations, red for high *k*_cat_ mutations. In (c) and (d), the high *k*_cat_ mutations and low *k*_cat_ mutations were used to represent mutations with higher and lower experimentally-measured *k*_cat_ values when compared to the wild-type enzyme

To further evaluate the generalization ability of DeepEnzyme in predicting *k*_cat_ values for enzyme variants from the saturation mutagenesis experiment, we utilized our model to predict *k*_cat_ values for large-scale mutants of phosphate-irrepressible alkaline phosphatase of Flavobacterium (PafA)^38^ and rigorously examined the results based on experimental evidences. We firstly categorized all PafA mutations into two distinct categories: high *k*_cat_ mutations and low *k*_cat_ mutations, representing mutations with higher and lower experimentally-measured *k*_cat_ values compared to the wild-type enzyme (Fig. 4d), respectively. It indicates that our model may help to characterize the increased *k*_cat_ levels for mutations with higher experimentally-measured *k*_cat_ values, though not very obviously (Fig. 4c). The median predicted *k*_cat_ values for the high *k*_cat_ mutations were observed to be 15% higher than those for the low *k*_cat_ mutation (*P* value = 0.0033). These results underscore the potential of DeepEnzyme in predicting the *k*_cat_ for enzyme variants from the saturation mutagenesis experiment.

### DeepEnzyme could identify key residue sites impacting enzyme activity

The active site of an enzyme, characterized by specific amino acid residues in the protein structure, is pivotal for enzyme catalysis. Similarly, the formation of enzyme-substrate complexes necessitates the presence of specific binding sites in the protein structure in order for the enzyme to interact with the substrate^39^. In this context, we sought to investigate whether DeepEnzyme possesses the capability to identify the importance of active sites and binding sites within a protein 3D structure. As the first example, we examined the case of PafA mentioned earlier. Leveraging site annotation data from the UniProt database, all residue sites in PafA were categorized into “binding/active sites” and “general sites”. The structure vectors were computed by DeepEnzyme, followed by normalization using min-max normalization to obtain the weight score for each residue site. In addition to PafA, P00558 in the human EMP pathway was also used to validate DeepEnzyme performance. The results clearly displayed that the weight scores of binding/active sites in PafA and P00558 were notably higher than those of the general sites (Fig. 5a, Fig. 5b, Fig. 5d, and Fig. 5e). At the same time, high-weight sites (These 2% residue sites with the highest weight scores) and binding/active sites in proteins are adjacent in structural space (Fig. 5c) or even have overlapping parts (Fig. 5f). Thus, DeepEnzyme showcased its potential in distinguishing and highlighting the biologically significant regions within protein 3D structures, which may provide valuable clues for rational engineering of enzymes.

**Fig. 5.**
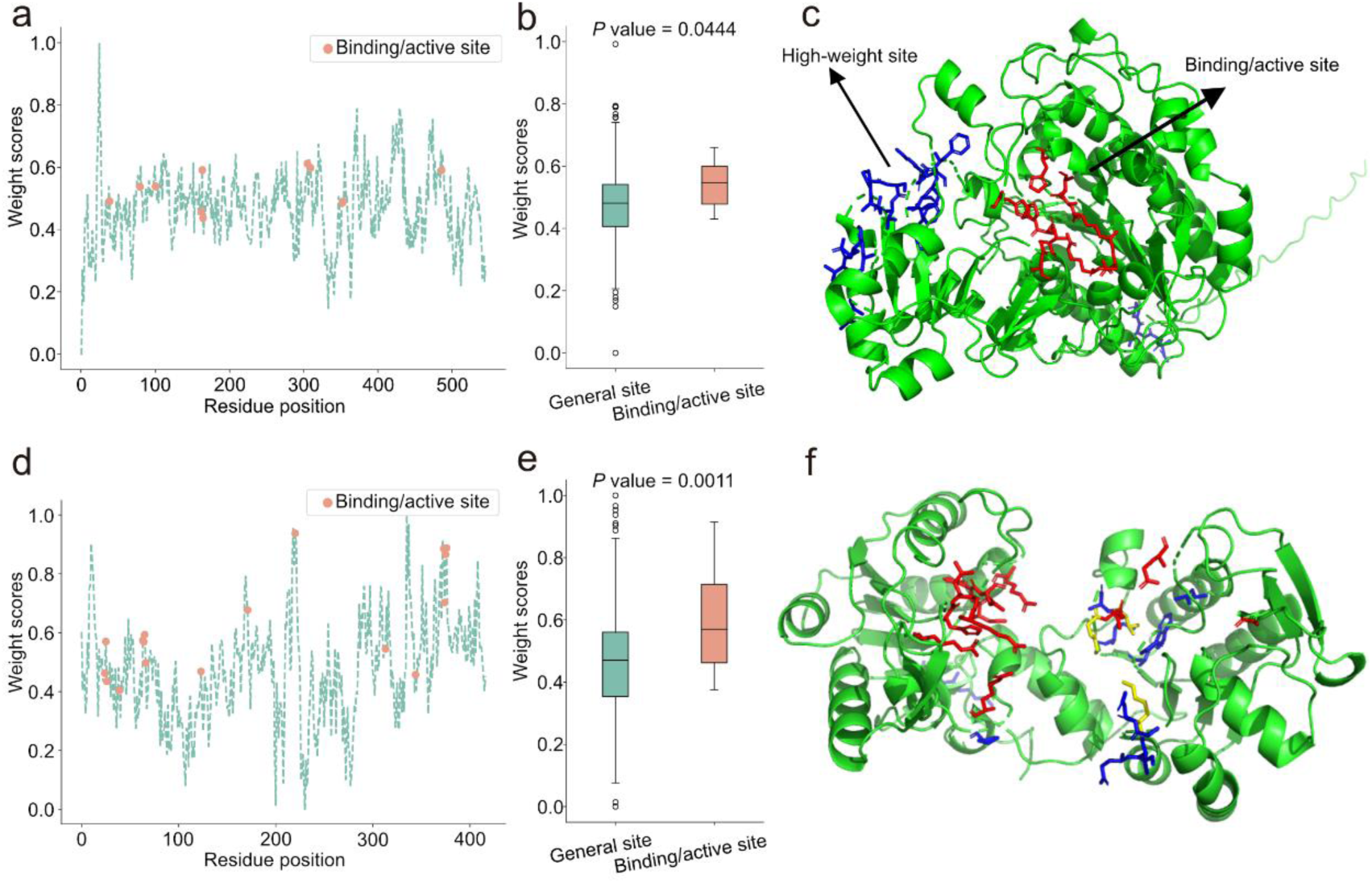
Comparison between the binding/active site and high-weight site (These 2% residues sites with the highest weight scores calculated by DeepEnzyme) within protein 3D structures. (a) The weight scores of different residue sites in PafA; the red points are binding/active sites. (b) Comparison of the weight scores between the binding/active sites and general sites in PafA; the green box line indicates general sites, and the red box line indicates binding/active sites. (c) The distribution of the binding/active sites and high-weight sites regions within the 3D structure of PafA, the red region represents for binding/active sites, the blue region for high-weight sites. (d) The weights of different residue sites in P00558; the red points are binding/active sites. (e) Comparison of the weights between the binding/active sites and general sites in P00558, where the green box line indicates general sites, and the red box line indicates binding/active sites. (f) The distribution of the binding/active sites and high-weight sites regions within the 3D structure of P00558, the red region represents for binding/active sites, the blue region for high-weight sites, the yellow region for the overlap between the two.

### Application cases: predict the *k*_cat_ for enzymes from genome-scale metabolic models

Genome-scale metabolic models (GEMs) are invaluable tools for characterizing cellular metabolism, often encompassing thousands of enzymes that catalyze various reactions. However, determining experimentally *k*_cat_ for a significant portion of enzymes in GEMs remains a challenging task. In this regard, DeepEnzyme presents itself as a convenient toolbox for predicting enzyme *k*_cat_ at the genome-scale. To showcase the potential applications of our model, we gathered and processed GEMs from diverse organisms, including *E. coli*, *M. musculus*, *S. cerevisiae*, and *H. sapiens*^40^, before *k*_cat_ prediction. Leveraging substrate and enzyme information, the genome-scale *k*_cat_ values could be predicted automatically (Fig. 6a and Fig. 6b). It shows that, at a holistic level, all *k*_cat_ values from multiple organisms or a single one exhibited a normal distribution, reflecting the obvious variance in predicted *k*_cat_ from different organisms.

**Fig. 6.**
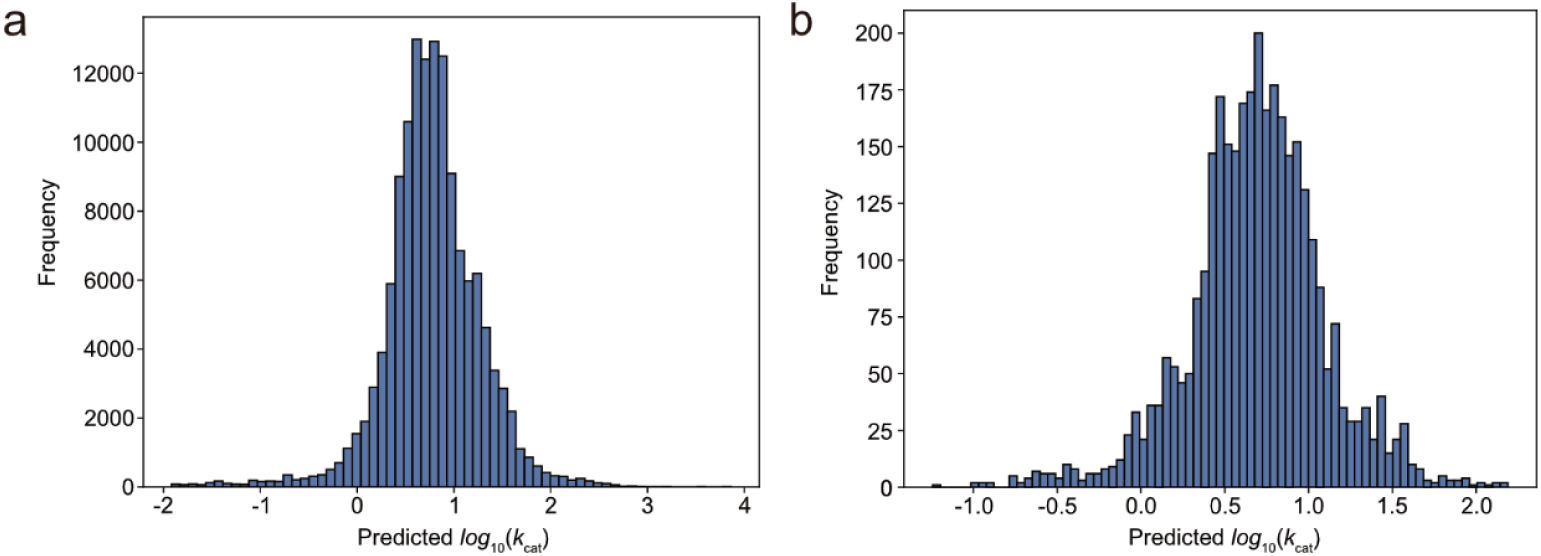
Predicted *k*_cat_ values for enzyme-catalyzed reactions in genome-scale metabolic models. (a) Distribution of *k*_cat_ values predicted by our model for enzyme-catalyzed reactions in metabolic models including those for *Homo sapiens*, *Mus musculus*, *Saccharomyces cerevisiae*, and *E*. *coli*^40^. (b) Distribution of *k*_cat_ values predicted by our model for enzyme-catalyzed reactions from the GEMs of *Geobacter metallireducens* GS-15 (BiGG ID: iAF987).

## Discussion

The turnover number is a fundamental parameter for understanding metabolism, growth, and resource allocation^4,5^. However, experimentally determining turnover numbers can be a time-consuming and labor-intensive process. Herein, we present DeepEnzyme, an enhanced model to improve enzyme *k*_cat_ prediction by utilizing advanced deep learning architectures such as Transformer and GCN (Fig.1). DeepEnzyme skillfully extracts and aggregates essential features from substrate, protein 1D sequences and 3D structures. With significantly enhanced prediction accuracy, DeepEnzyme surpasses previously reported models in *k*_cat_ prediction. Specifically, it achieves an impressively higher R^2^ value at about 0.6 on the test dataset (Fig.2 and Fig.3), outperforming both TurNuP (R^2^=0.40)^14^ and DLKcat (R^2^=0.43)^13^. PreKcat^41^, which utilizes the latest pre-trained large model for *k*_cat_ prediction, was released when we prepared this manuscript. However, it is difficult to directly compare the prediction performances of DeepEnzyme with PreKcat as the source code for training PreKcat was not accessible. According to Yu^41^, PreKcat could outperform DeepEnzyme in prediction accuracy due to its usage of the SMILES Transformer model and the pre-trained protein language model. However, with fewer parameters and smaller model size, DeepEnzyme is more than 2,000 times faster than PreKcat when completing the same prediction task, i.e., predicting *k*_cat_ for 7630 enzyme variants, assuming that the protein 3D structures are ready in advance (testing with the PreKcat code example on DELL Precision 5820 Tower).

DeepEnzyme shows robustness in predicting *k*_cat_ values for remote enzyme homologs. The degree of similarity in protein sequences originated from the test and the training datasets could significantly influence the performance of deep learning models, and the prediction accuracy would decrease as the sequence dissimilarity increases^13,14^. Through careful preprocessing of the sequence dataset, we significantly reduce the tendency of DeepEnzyme in overfitting during training, thereby improving the generalization ability of the model and ensuring the reliable *k*_cat_ predictions across diverse levels of sequence similarity. By comparison, even for enzymes with smaller sequence similarity (0–40%) compared to that from the training dataset, DeepEnzyme still exhibits outstanding prediction accuracy with an R^2^ value of 0.54 (Fig.3), thus successfully outperforming previous models^13,14^. It is well known that protein 3D structures are more evolutionarily conservative than amino acid sequences. Thus, extracting and utilizing the enzyme’s structural features during our model training allowed for a considerable improvement in the accuracy of k*_cat_* prediction regardless of variation in the protein sequence similarity.

Furthermore, DeepEnzyme could help to evaluate of the influence of saturation mutagenesis on enzyme catalytic efficiency. In this aspect, the performance of DeepEnzyme was demonstrated using two enzymes, PafA^38^ and CYP2C9^37^, for which abundant experimental data on point mutations are available. The results indicate that our model can, to a considerable extent, reflect how single mutations affect enzyme catalytic efficiency (Fig.4). Moreover, DeepEnzyme assigns higher weight scores to active sites and binding sites within a protein 3D structure. As these functional sites are closely correlated to enzyme function and activity^14^ (Fig.5), our model potentially help to establish the mapping from sequences, to 3D structures, until to protein functions. So, the *k*_cat_ prediction together with the weight score inference for each residue site by DeepEnzyme could offer mechanical insights underlying the variance of enzyme catalytic efficiency, which may empower the rational design of more efficient enzymes for experimental test and validation.

While DeepEnzyme outperforms the existing public models in terms of *k*_cat_ prediction accuracy, it could be further enhanced in the following aspects. Firstly, the dataset used in this work encompassed less than 12,000 enzyme-substrate pairs, which is relatively smaller compared to the larger training datasets widely used in deep learning models. Expanding the dataset to encompass a more diverse enzyme-substrate pairs could further enhance the predictive performance of DeepEnzyme. Secondly, the pivotal experimental parameters such as pH and temperature were not taken into account during the model training in this work, which may contribute to discrepancies between predicted and measured *k*_cat_ values for certain enzymes. To further enhance the performance of DeepEnzyme, more standardized experimental datasets under diverse environmental conditions should be considered and collected. Additionally, multiple advanced pre-training models can be further merged into the current framework of DeepEnzyme to improve the prediction accuracy. It is envisioned that, together with other deep learning models, DeepEnzyme will undoubtedly provide new chances to characterize the internal correlations among 1D sequence, 3D structures and enzyme catalytic efficiency, thus accelerating the protein engineering in the coming years.

## Methods

### Data preprocessing

In the reconstruction of DeepEnzyme, the DLKcat dataset was first downloaded^13^, which is openly accessible and contains details about the enzyme and substrate, including the EC number, organism, enzyme sequence, simplified molecular-input line-entry system (SMILES), and *k*_cat_ value. In order to mitigate the impact of high sequence similarity among enzyme-substrate pairs in the dataset during model training, a preprocessing step was applied to retain only the enzyme-substrate pairs with the highest enzyme sequence length among those with identical substrates and sequence similarity greater than 90%. In order to assess how comparable the enzyme sequence was across the dataset, the MMseqs2^31^ approach was employed. Only the enzyme-substrate pairs with the longest enzyme sequences were retained, with the aim of reducing bias and overfitting that may result from highly similar sequences. This preprocessing strategy allows the model to focus on more diverse and representative enzyme-substrate pairs, ultimately enhancing its generalization ability and predictive performance. After data preprocessing, the dataset includes 6,496 unique protein sequences and a total of 11,927 distinct combinations of enzymes and substrates. Additionally, the dataset is randomly divided into groups of training, validation, and testing with ratio at 80%, 10%, and 10% for the development of DeepEnzyme.

### Enzyme structure prediction and contact map generation

In order to predict the enzyme structure and gather the structural data of enzymes, ColabFold^22^ was used to predict protein structure, which is approximately 40-60 times faster than AlphaFold2. As a whole, the predicted structures have an average predicted local distance difference test (pLDDT) score of 92.67, with over 98% of the enzymes having a pLDDT score higher than 75 (Fig. 1), indicating that the structures used in this work are of high quality. Next, the protein 3D structure was converted into a contact map, which could transform the protein structures into a reasonable format suitable for deep learning algorithms. For creating the contact map, the residues within the protein 3D structures were defined as nodes, while two nodes were connected by a linker if the distance between the respective *C_α_* atoms of two residues was smaller than10Å. It has been demonstrated that this procedure could successfully capture crucial structural features used in previous deep learning models for protein function prediction^10,42^.

### Deep learning pipeline for turnover number prediction in DeepEnzyme

To accurately predict enzyme *k*_cat_, an end-to-end deep learning model that combines the Transformer and Graph Convolutional Network (GCN) architectures has been conceived and developed. The Transformer and GCN are particularly designed to extract hidden features from both protein 1D-sequence and 3D structures. Here, the thorough description of the pipeline for training and testing DeepEnzme is given as following.

#### The protein sequence baseline

In order to extract protein sequence features, the Transformer model, which has excellent performances in natural language processing, is employed in this work. Similar to DLKcat, the protein sequence is split into an overlapping sequence composed by n-grams of amino acids. Before the Transformer, an embedding layer is employed to convert the protein sequence into a word tensor. The hidden vectors of the protein sequence are then determined using word embedding as input, followed by a Transformer encoder with position encodings (Eq.1), 4 multi-head attention layers (Eq.2,3), and a hidden dimension ( *d_model_* ) of 64.

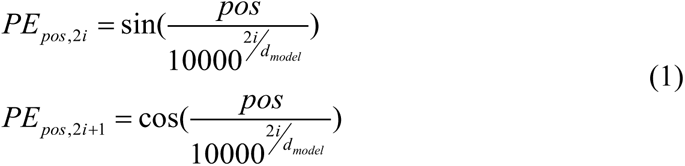

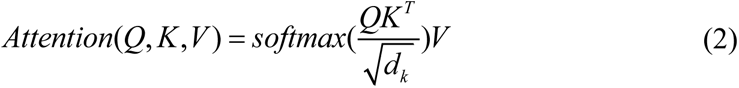

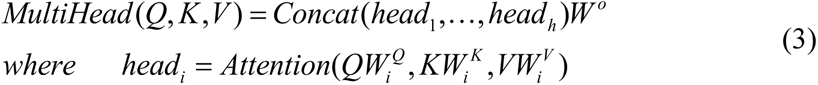

Where pos is the position of the amino acid, d_model_ is the dimension of the vector, Q, K, V stand for Query vector, Key vector and Value vector respectively.

#### The protein and substrate structure baseline

The contact map is used to represent the protein structure, and the substrate graph from the DLKcat database is used to characterize substrate information. GCN (Eq.4) generates substrate vectors and protein structure vectors as its final output via a convolutional neural network.

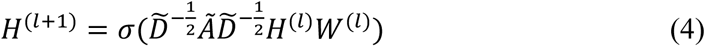

*The neural attention mechanism baseline.* To obtain the final *k*_cat_ value output, the neural attention mechanism is incorporated, which enables us to capture the attention weights of various residue sites in the enzyme^43^. This attention mechanism pays more attention into the important fragments of the protein sequence and structure. This attention mechanism (Eq.5-7) works in conjunction with the Transformer and GCN components, which makes it feasible to extract features from the protein sequence and 3D structures.

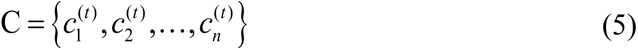

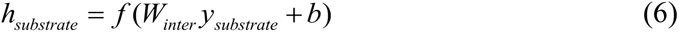

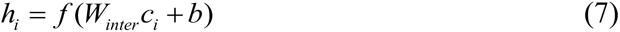

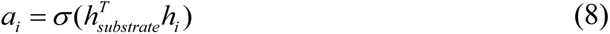

where *C* is a set of hidden vectors representing the protein sequence, 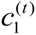 to 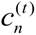 are sub-hidden vectors for split subsequences within the protein sequence, *y_substrate_* is the molecular vector of the substrate, *W* and *b* are the weight matrix and the bias vector, respectively, used in the neural network, *f* is a nonlinear activation function, for example, ReLU (Rectified Linear Unit)^44^, *α_i_* is the final attention weight value, which is calculated using the attention mechanism, *σ* is the element-wise sigmoid function, used for specific calculations within the model, and *T* represents the transpose function.

#### Evaluation metrics

In this study, three performance metrics are used to evaluate the prediction performances of the model: the coefficient of determination (R^2^, Eq.9) the root mean square error (RMSE, Eq.10), and the Pearson correlation coefficient (PCC, Eq.11). These metrics are widely used in regression analysis to examine the correlation and accuracy of predictions against experimental values.

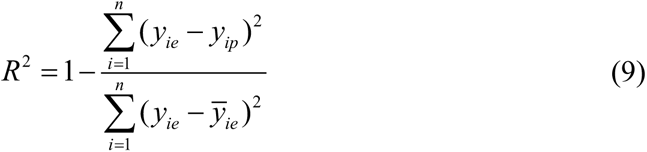

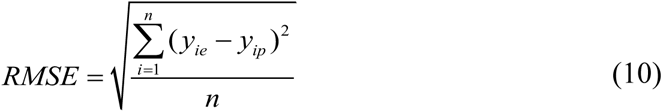

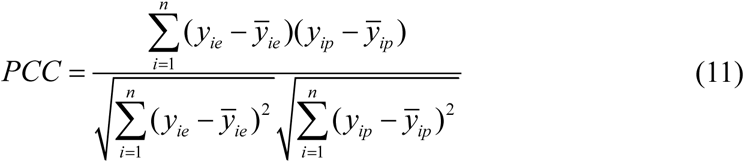

*Where y_ie_* represents the experimental value of *k*_cat_, *y_ip_* represents a predicted value for *k*_cat_, and *ȳ* represents the corresponding averages.

### Comparison of different deep learning models in *k*_cat_ prediction

Utilizing the code and datasets stored in GitHub by Li et al.^13^ and Kroll et al.^14^, the prediction performance of DLKcat and TurNuP was evaluated respectively. Based on the similarity of the protein sequences, the DLKcat test dataset, the TurNuP test dataset, and our test dataset were separated into three distinct subsets. MMseqs2^31^ was utilized to determine the degree of similarity between the protein sequences in the training dataset and the test dataset, FoldSeek^32^ and US-align^33^ were used to determine the degree of partial protein structural similarity. The R^2^ value calculated from the test dataset was used to evaluate the prediction accuracy of above different deep learning models.

### Evaluative the prediction performance of DeepEnzyme based on large-scale enzyme saturation **mutagenesis** experiments

Sequence data for CYP2C9 variants and PafA mutations were obtained from the studies of Amorosi, C.J., et al.^37^, and Markin, C.J., et al.^38^, respectively, and ColabFold was used to predict the 3D structures for all variants of CYP2C9 and PafA in order to generate the corresponding independent datasets containing substrate, protein sequence, and protein structures. The *k*_cat_ values were predicted directly on the two datasets with DeepEnzyme, and the predictions were compared in groups, which were defined by Amorosi, C.J., et al.^37^ or the *k*_cat_ values in the study by Markin, C.J., et al.^38^ .

### Interpretation analysis of DeepEnzyme in ranking the key residue sites related to enzyme activities

Wild-type PafA and P00558 were used as input for the *k*_cat_ prediction by DeepEnzyme, and the data in each column of the protein structural feature matrix extracted by GCN were averaged, while min-max normalization was performed to obtain the weight of each residue site, and the sites were grouped according to the site classification (binding/active sites and general sites) in the UniProt database^8^. Then, the weight score of sites was compared in groups.

### Statistical analysis

In all statistical analysis, the two-sided t-test in the Python package SciPy was used^45^.

## Data availability

The large file used to train the model could be found in: https://figshare.com/authors/Leo_wang/12865928.

## Code availability

The codes are accessible at https://GitHub.com/hongzhonglu/DeepEnzyme.

## Author contributions

HZL designed the research. TW performed the research. HZL and TW analyzed the data. All authors interpreted the results, discussed, drafted and approved the final manuscript.

## Acknowledgements

This work is supported by grant 2022YFA0913000 from the National Key R&D Program of China, Shanghai Pujiang Program, grant 22208211 and 22378263 from the National Natural Science Foundation of China (NSFC).

